# Lactate dehydrogenase-induced DNA Topoisomerase 1 is a novel regulator of smooth muscle cell proliferation and remodeling in pulmonary arterial hypertension

**DOI:** 10.1101/2025.11.10.687749

**Authors:** Lifeng Jiang, Iryna Zhyvylo, Dmitry Goncharov, Tapan Dey, Neil J. Kelly, Leyla Teos, Lisa Franzi, Aisha Saiyed, Nicholas J. Kenyon, John R. Greenland, Paul J. Wolters, Stephen Y. Chan, Horace Delisser, Tatiana V. Kudryashova, Elena A. Goncharova

**Affiliations:** University of California, Davis School of Medicine, Davis, CA, USA; Veterans Affairs Pittsburgh Healthcare System, Pittsburgh, PA, USA; Center for Pulmonary Vascular Biology and Medicine, Pittsburgh, Heart, Lung, and Blood Vascular Medicine Institute, University of Pittsburgh, Pittsburgh, PA, USA; Division of Cardiology, University of Pittsburgh School of Medicine, Pittsburgh, PA, USA; VA Northern California Healthcare System, Mather, CA; University of California, San Francisco School of Medicine; San Francisco Veterans Affairs Health Care System, San Francisco, CA, USA; University of Pennsylvania Perelman School of Medicine, Philadelphia, PA, USA

**Keywords:** lactylation, non-histone proteins, apoptosis, pulmonary vascular disease, TOP1, Akt, mTOR

## Abstract

Pulmonary arterial hypertension (PAH) manifests by increased proliferation and survival of pulmonary vascular cells in small pulmonary arteries (PAs), PA remodeling and unresolved increase of PA pressure. PA smooth muscle cells (PASMCs) in PAH undergo metabolic shift to glycolysis resulting in over-production of lactate, hyper-proliferation, and apoptosis resistance, but the mechanisms are not completely understood. By using lung tissues and pulmonary vascular cells from PAH and non-diseased human lungs, unbiased proteomics, network analysis, and gain-and-loss of function approaches, we here report that up-regulation of lactate dehydrogenase A (LDHA)-lactate axis promotes PASMC-specific over-lactylation and consequent over-accumulation of DNA topoisomerase 1 (TOP1) in small remodeled PAs from PAH lungs, leading to the up-regulation of Akt-mechanistic target of rapamycin 1 (mTORC1) signaling, hyper-proliferation, and reduced apoptosis. Smooth muscle-specific LDHA knockdown prevented, and Ldha inhibitor oxamate reversed SU5416/hypoxia-induced TOP1 accumulation, pulmonary vascular remodeling, and pulmonary hypertension (PH) in mice. Pharmacological inhibition of TOP1 with indotecan suppressed Akt-mTORC1, decreased proliferation, induced apoptosis in human PAH, but not control PASMCs, and reversed PA remodeling, PH, and RV dysfunction in rats. Collectively, these data provide a novel mechanistic link from LDHA-driven lactate over-production through lactylation and overaccumulation of TOP1, to the up-regulation of Akt-mTORC1, hyper-proliferation and apoptosis resistance of PASMCs, pulmonary vascular remodeling, and PH, and identify TOP1 as a new potentially attractive molecular target for the remodeling-focused therapeutic intervention.

**Take-home message:** LDHA-lactate-induced over-lactylation and overaccumulation of Topoisomerase 1 (TOP1) promotes pulmonary artery smooth muscle cell hyper-proliferation, remodeling, and pulmonary arterial hypertension, which are reversed by TOP1 inhibitor indotecan.

## INTRODUCTION

Pulmonary arterial hypertension (PAH) is a progressive and rapidly fatal disease with a poor prognosis and limited treatment options (1–5). PAH manifests by vasoconstriction and remodeling of small pulmonary arteries (PAs), leading to increased PA pressure (PAP), elevated right ventricular (RV) afterload, and premature death of heart failure (6). Despite recent advances, available therapies fail to reverse pulmonary vascular remodeling or prevent disease progression (7–10). Increased proliferation and survival of resident PA cells, supported by metabolic shift to glycolysis, play an important role in the remodeling of small muscularized PAs and overall PAH (11, 12), but the mechanistic link between dysregulated metabolism and PA remodeling is not completely understood.

Lysine lactylation (Kla) is a post-translational modification of histone and non-histone proteins. Kla is mediated, at least in part, by the availability of lactate, a byproduct of glycolysis generated from pyruvate by lactate dehydrogenase A (LDHA). High serum LDH correlates with severity and mortality in idiopathic PAH (13). PA smooth muscle cells (PASMCs) in PAH switch to the glycolytic energy generation, leading to elevated lactate production and secretion (11, 14–17). LDHA and/or lactate could promote PASMC proliferation (18–21), while LDHA inhibition attenuates experimental pulmonary hypertension (PH) (18). Dysregulated lactylation plays important role in cancer and cardiovascular diseases (22, 23), and lactylation of histones contributes to the hypoxia-induced experimental PH (24). The role of protein lactylation in PAH remains to be established.

The goal of this study was to determine the status and mechanistic consequences of protein lactylation in PAH pulmonary vasculature and test potential attractiveness of this signaling as a target pathway for remodeling-focused therapeutic interventions to treat PAH. We report that LDHA-lactate-dependent over-lactylation and consequent overaccumulation of DNA topoisomerase 1 (TOP1) promotes activation of Akt-mammalian target of rapamycin complex 1 (mTORC1) axis, hyper-proliferation and apoptosis resistance of PAH PASMCs, pulmonary vascular remodeling, and PH. These data support TOP1 as a molecular target to reduce PASMC hyper-proliferation and reverse established PH.

## RESULTS

### TOP1 is over-lactylated and over-accumulated in PASMCs in human PAH lungs

To determine the status of global protein lactylation (Kla) in PAH pulmonary vasculature, we performed immunohistochemical and immunoblot analyses of lung tissue sections and isolated pulmonary vascular cells from human non-diseased (control) and PAH lungs (Table S1) with anti-pan-Kla antibody. We detected markedly higher Kla levels in smooth muscle α-actin (SMA)-positive areas of small remodeled PAs (Figure 1A) and in early-passage PASMCs from human PAH lungs, which exhibited significant unstimulated proliferation (Figures 1B-D). We detected no differences in Kla levels in PA endothelial cells (PAECs) or PA adventitial fibroblasts (PAAFs) from human PAH lungs compared to controls (Figure S1), indicative of PASMC-specific increase in global protein lactylation in PAH PAs.

**Figure 1.**
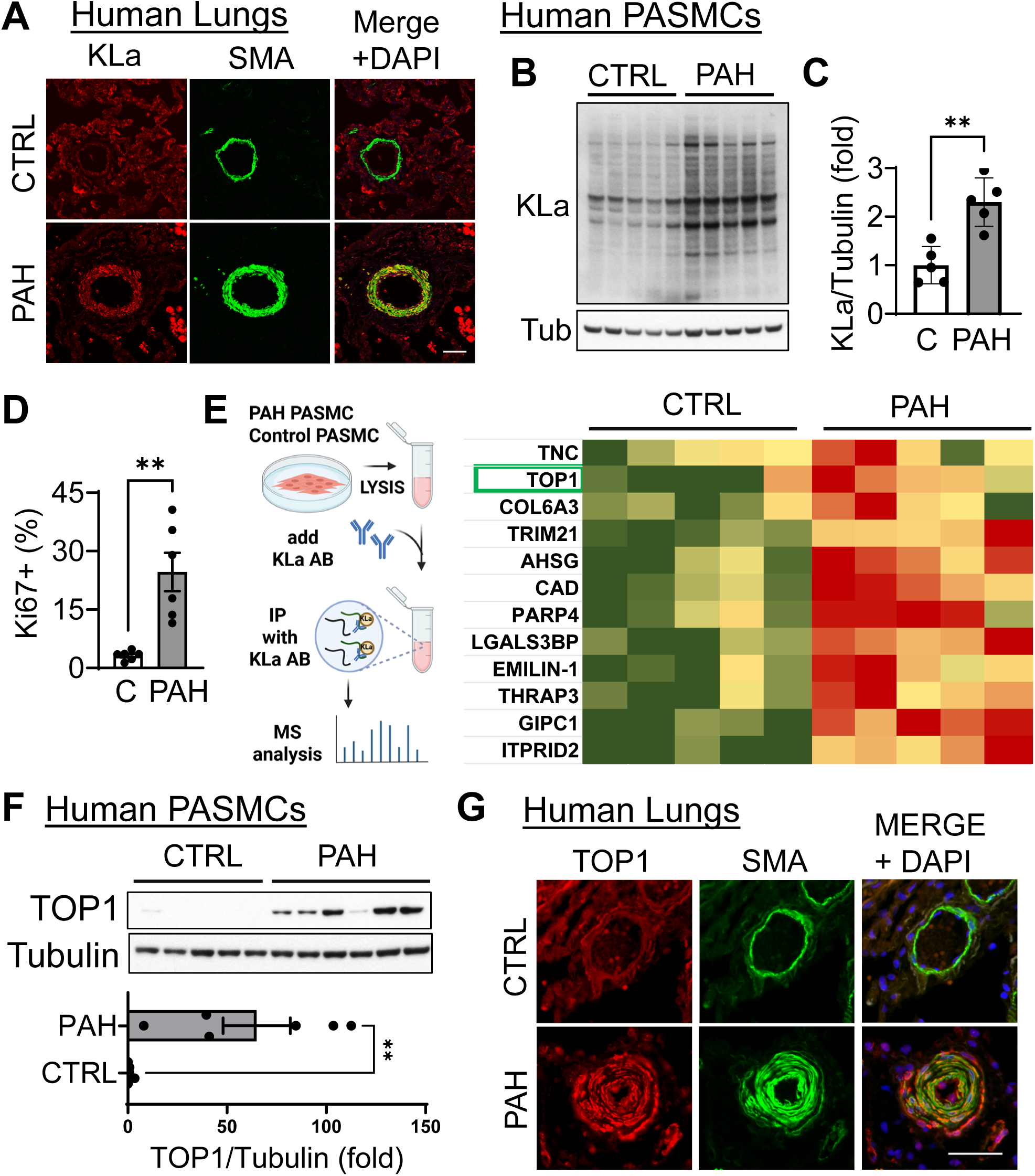
TOP1 is over-lactylated and over-accumulated in PASMC from human PAH lungs. **A:** Immunohistochemical analysis of human lungs to detect pan-L-lactyllysin (Kla). Images are representative of 5 subjects/group. Red -Kla, green -SMA, blue -DAPI. Bar=50 µm. **B-D:** Human control and PAH PASMCs were subjected to immunoblot (n=5 subjects/group) (**B, C**) and proliferation (Ki67, n=6 subjects/group) (**D**) analyses. Data are means±SE, **p<0.01 by Mann Whitney U test. **E**: Significantly differentially lactylated proteins identified by proteomic analysis of anti-Kla antibody immunoprecipitants from whole cell lysates of non-diseased (control) and PAH PASMCs (5 subjects/group). **F**: Immunoblot analysis of human control and PAH PASMCs to detect indicated proteins. Data are means±SE, n=5 subjects/control, 6 subjects/PAH group, **p<0.01 by Mann Whitney U test. **G:** IHC analysis of human lung tissues. Red -TOP1, green -SMA, blue -DAPI. Bar=50 µm. Representative images from 3 subjects/group.

Mass spectrometry analysis of anti-Kla antibody immunoprecipitants identified 12 overactivated non-histone proteins in human PAH PASMCs compared to controls (Figure 1E). Four proteins (TOP1, PARP4, LGALS3BP, GIPC1), inhibitors of which are in clinical use or preclinical development as anti-proliferative drugs, were selected for further validation. Out of four analyzed proteins, only TOP1 was significantly over-accumulated in human PAH PASMCs compared to controls (Figures 1F, S2). Further supporting our observations, TOP1 protein levels were markedly increased in SMA-positive areas of small remodeled PAs in human PAH lungs (Figure 1G). Together, this data demonstrate that TOP1 is hyper-lactylated and over-accumulated in PASMCs from human PAH lungs.

### TOP1 overaccumulation in human PASMCs is driven by LDHA-lactate

Because protein lactylation is modulated, at least in part, by lactate availability, we reasoned that TOP1 over-lactylation and over-accumulation could be caused by elevated lactate and/or LDHA levels. Supporting this idea, immunohistochemical and immunoblot analyses revealed marked increase in LDHA protein levels in SMA-positive areas of small muscular PAs from human PAH lungs (Figure 2A-C). Isolated human PAH PASMCs had significantly higher LDHA mRNA and protein content, and elevated lactate secretion compared to non-diseased controls (Figure 2D-G). We detected no differences in LDHA protein levels between control and PAH PAECs and PAAFs (Figure S3). These data show that PASMCs from subjects with PAH have up-regulated LDHA and lactate over-production.

**Figure 2.**
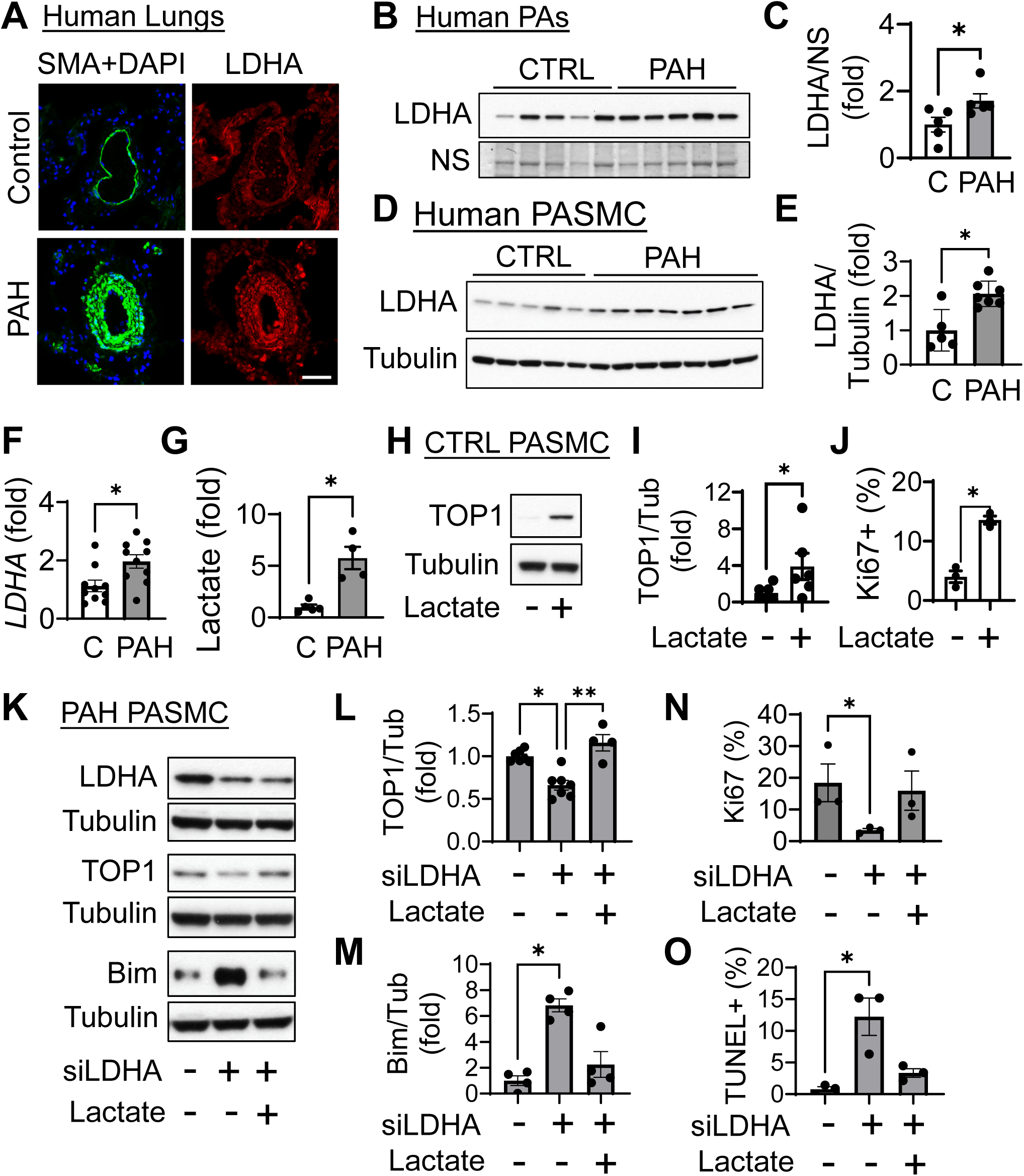
Up-regulation of LDHA-lactate is required for TOP1 over-accumulation, hyper-proliferation and survival of human PAH PASMCs. **A:** Immunohistochemical analysis of human lung tissues to detect LDHA (red), SMA (green), and DAPI (blue); representative from 5 subjects/group. Bar=50µm. **B, C:** Immunoblot analysis of small muscular PAs (<1.5 mm outer diameter) from human PAH and control lungs to detect LDHA protein levels. **D-G**: Immunoblot (**D,E**), RT-PCR (**F**), and intracellular lactate levels (normalized by cell numbers) (**G**) of human control and PAH PASMCs. **B-G:** Data are means±SE; n=5-10 subjects/group, *p<0.05, **p<0.01 by Mann Whitney U. **H-J:** Human control PASMC were treated with 10mM lactate or diluent (-) for 48 hours. Then, immunoblot analysis was performed (n=6 subjects/group) (**H, I**), proliferation was measured using Ki67 assay (n=3 subjects/group) (**J**). Representative images are shown; Data are means±SE; *p<0.05 by Mann Whitney U test. **K-O**: Immunoblot (**K-M**), proliferation (Ki67) (**N**), and apoptosis (*In Situ* Cell Death Detection Kit) analyses (**O**) of human PAH PASMC transfected with siRNA LDHA or control siRNA GLO and treated with 10mM lactate or diluent (-) for 48 hours. Data are means±SE, n=4-7 subjects/group; *p<0.05, **p<0.01 by Kruskal-Wallis (Dunn).

Treatment with lactate induced TOP1 protein accumulation and proliferation of control human PASMCs (Figure 2H-J). In contrast, siRNA-induced depletion of LDHA in human PAH PASMCs reduced TOP1 protein levels and cell proliferation, while increasing accumulation of pro-apoptotic Bim and promoting apoptosis. All these effects were reversed by lactate supplementation (Figure 2K-O). In aggregate, these data show that TOP1 over-accumulation, hyper-proliferation, and apoptosis resistance of human PAH PASMCs are induced by LDHA through lactate over-production.

### LDHA promotes TOP1 overaccumulation, pulmonary vascular remodeling, and pulmonary hypertension in mice

To evaluate the relationship between LDHA and TOP1 signaling *in vivo*, we developed mice with SM-specific Ldha depletion (Ldha-/-^SMC^) by crossing SM22-Cre mice and Ldha fl/fl mice (Figure S4), and induced PH by SU5416/hypoxia (SuHx) exposure (25, 26) (Figure 3A). Three weeks after PH induction, wild-type (WT), but not Ldha-/-^SMC^ mice, developed significant increase in TOP1 protein content in SMA-positive areas of small muscular PAs, PA medial thickness (PA MT), and systolic right ventricular pressure (sRVP) (Figure 3B-F). We detected no differences in systolic left ventricular pressure (sLVP) among groups (Figure S5). These data show that smooth muscle Ldha is required to induce TOP1 overaccumulation, pulmonary vascular remodeling, and PH in mice.

**Figure 3.**
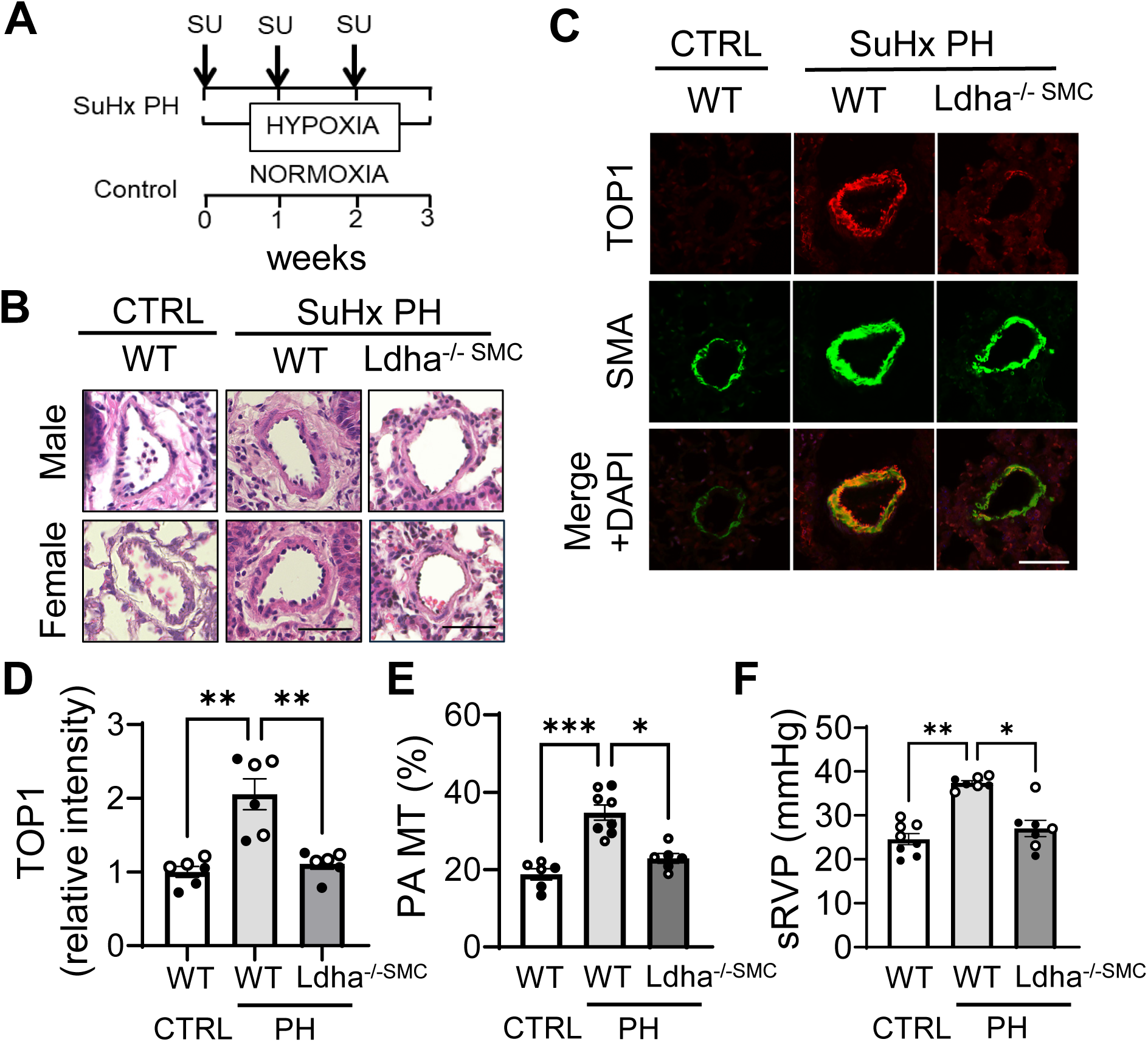
Smooth muscle Ldha is required for TOP1 up-regulation, pulmonary vascular remodeling, and PH in mice. **A**: Male (○) and female (●) wild type (WT) and Ldha-/-^SMC^ (-/-) mice were exposed for 21 days to normobaric hypoxia (10% O_2_) and received SU5416 injections (s.q., 20 mg/kg) on days 1, 8, and 15 of hypoxia exposure. Control group was same-sex same-age WT mice maintained under normoxia. On day 22, biventricular hemodynamic analysis and lung tissue collection for immunohistochemical and morphologic analyses were performed. **B, C:** Representative hematoxylin & eosin (H&E) (**B**) and immunohistochemical images (**C**) of small PAs from minimum of 6 (minimum of 3 male and minimum of 3 female) mice/group are shown. Bar=50µm. Red -TOP1, green -SMA, blue -DAPI. **E-G**: Relative TOP1 levels in SMA-positive areas of small PAs calculated using optical density (OD) (**D**), PA medial thickness (**E**), and systolic right ventricular pressure (sRVP) (**F**) of indicated groups. Data are means±SE from 6-8 mice/group, *p<0.05, **p<0.01, ***p<0.005 by Kruskal-Wallis (Dunn).

To test whether LDHA supports TOP1 up-regulation in established PH, we used oxamate, a non-metabolizable isosteric form of pyruvate that serves as a competitive inhibitor of LDHA (27–29). After three weeks of PH induction by SuHx, when PH is already developed (25, 26), male WT mice, still kept under hypoxia, were randomly assigned to two groups and treated with oxamate or vehicle for two more weeks. Controls were same-age male mice maintained under normoxia (Figure 4A). Vehicle-treated mice showed significant increase in TOP1 in the SMA-positive areas of small muscular PAs, elevated PA MT, sRVP, and Fulton index (RV/[LV+S] ratio) compared to normoxia-maintained controls (Figure 4B-G), indicative of the SM-specific TOP1 overaccumulation, pulmonary vascular remodeling, PH, and RV hypertrophy. Oxamate normalized TOP1 levels, reduced remodeling, PH, and RV hypertrophy compared to vehicle-treated group (Figure 4B-G). We detected no differences in sLVP among groups (Figure 4H). Taken together, these data show that Ldha inhibition reverses SM-specific TOP1 overaccumulation and attenuates established PA remodeling and PH in mice.

**Figure 4.**
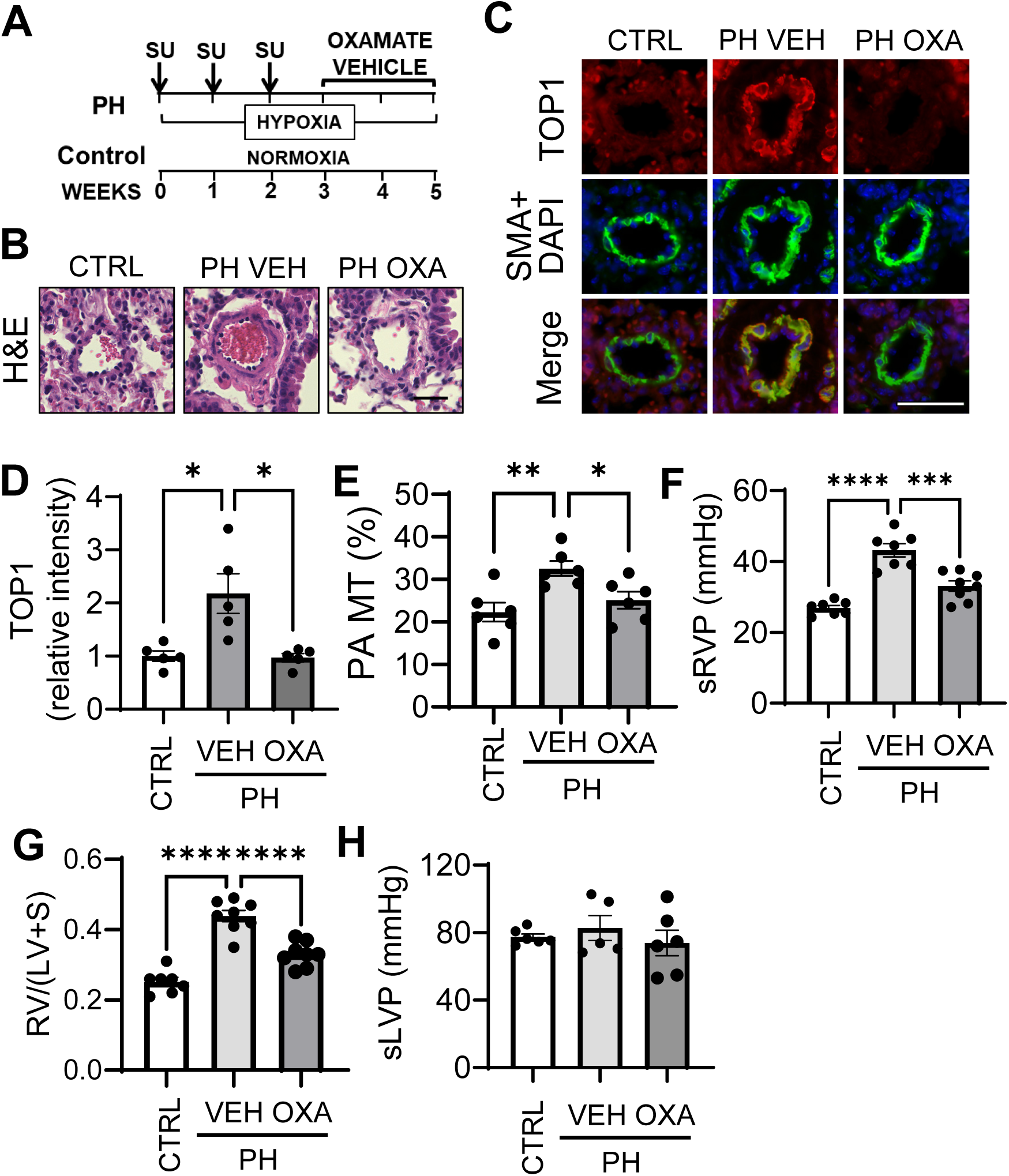
LDHA inhibitor oxamate reverses TOP1 overaccumulation, attenuates established PA remodeling and PH in mice. **A**: PH was induced in 12 months old male C57BL/6J mice by three weeks of hypoxia (10% O_2_) and three injections of SU5416 (s.q., 20 mg/kg) at days 1, 7, and 14. Oxamate (750mg/kg/day, i.p. (75) or vehicle were administrated at days 22-35. Controls were same-age male mice maintained under normoxia. **B, C:** H&E (**B**) and immunohistochemical analysis of lung tissue sections (**C**) of mouse lungs from indicated groups. Bar=50µm. Images are representative of 5-6 mice/group. **C**: Red -TOP1, green -SMA, blue-DAPI. **D-H**: TOP1 relative intensity (**D**) PA MT (**E**), sRVP (**F**), Fulton index (RV/(LV+S) (**G**), and sLVP (**H**) of indicated groups. **D, G:** 5-6 mice/group, 10 PAs/mouse. **F-H**: n=7-8 mice/group. Data are means±SE. *p<0.05, **p<0.01, ***p<0.005, ****p<0.0001 by Kruskal-Wallis (Dunn) (**E, F**) or ANOVA (Tukey) (**H-J**).

### TOP1 promotes proliferation and survival of human PAH PASMCs via Akt-mTORC1

To determine the mechanism(s) by which TOP1 promotes hyper-proliferation of human PAH PASMCs, we performed network analysis to predict signaling pathways relating TOP1 to the PAH interactome (26, 30–32) using previously identified PH genes (i.e. “PH Network”) (33) and gene set enrichment analysis. The key TOP1 interaction pathways were involved in regulating cell proliferation and apoptosis (Table S2). The top three highly scored pathways were Akt-mTOR, p53, and WNT-β catenin (Figure 5A, B, Table S2). To validate network analysis predictions, we transfected human PAH PASMCs with siRNA TOP1. We observed significant reduction of Akt-mTORC1 pathway assessed by S473-Akt and ribosomal protein S6 phosphorylation rates, a molecular signatures of Akt and mTORC1 activation (34), significantly reduced cell proliferation and elevated apoptosis compared to control siRNA-transfected cells (Figure 5C-G). siRNA LDHA, in turn, reduced S473-Akt and S6 phosphorylation rates in human PAH PASMCs (Figure 5 H-J), while treatment with lactate increased Akt and S6 phosphorylation in control PASMCs (Figure 5 K-M). siRNA TOP1 increased p53 protein levels in human PAH PASMCs without changes in β-catenin (Figure S6). Neither siRNA LDHA, nor lactate affected p53 protein content (Figure S7). Together, these data suggest that LDHA-lactate-driven TOP1 overaccumulation promotes human PAH PASMC hyper-proliferation and survival via up-regulating Akt-mTORC1.

**Figure 5.**
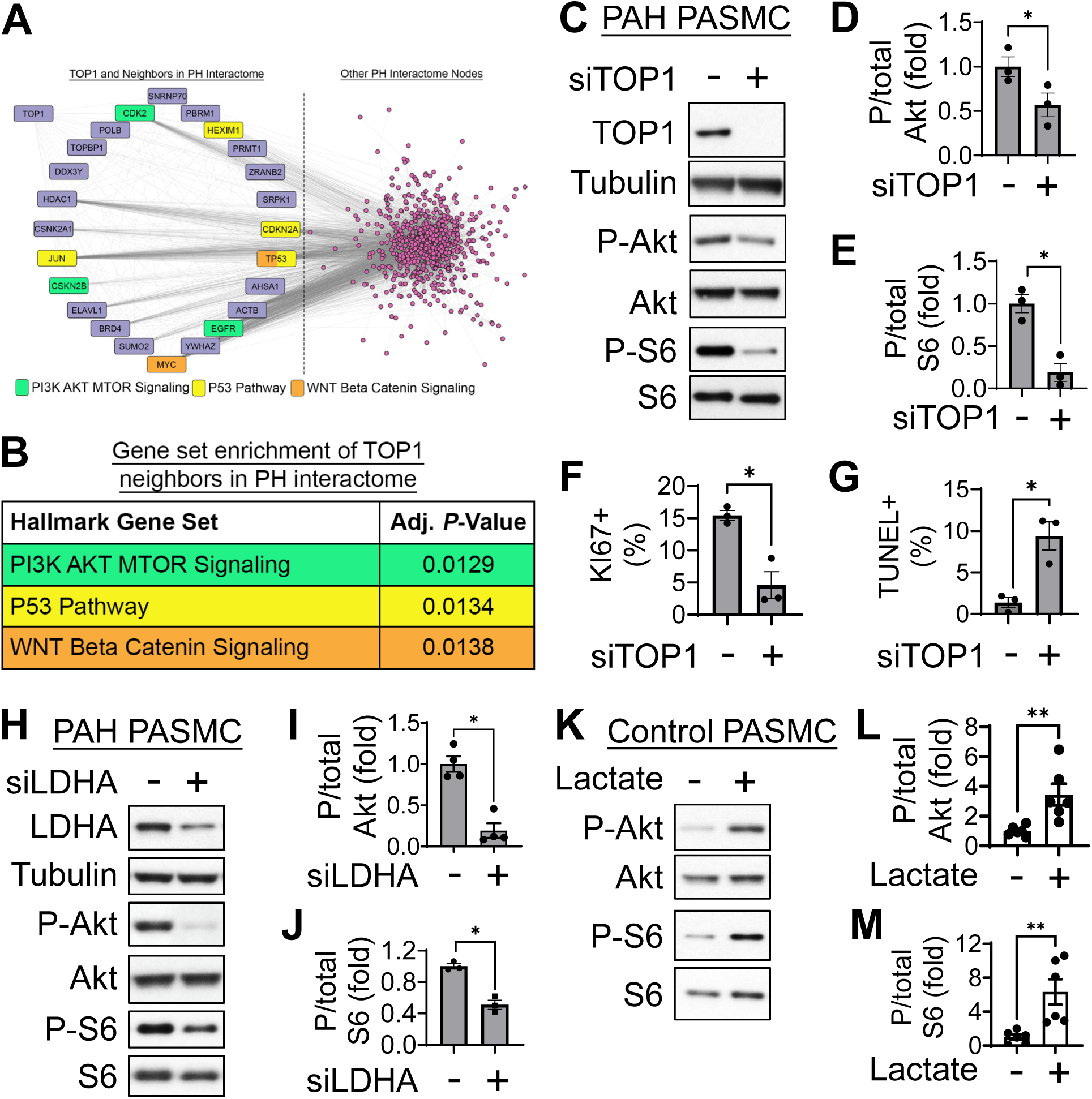
TOP1 promotes proliferation and survival of human PAH PASMCs via Akt-mTORC1. **A, B:** Network analysis (**A**) and gene set enrichment analysis (**B**) to predict the PH interactome as the largest connected component from the subset of the global interactome in which both interactors were PH-associated proteins inclusive of LDHA and TOP1. Hypergeometric *P*-values and were computed and adjusted for false discovery rate (FDR) by the Benjamini-Hochberg method using Python’s *scipy* module, version 1.13.0, default settings. **C-G**: Human PAH PASMCs were transfected with siRNA TOP1 or control siRNA GLO (-) for 48 hours followed by immunoblot analysis (**C-E**), proliferation (Ki67) (**F**), and apoptosis (*In Situ* Cell Death Detection Kit) assays (**G**). **H-M**: Immunoblot analysis of human PAH PASMCs transfected for 48 hours with siRNA LDHA or control si RNA GLO (-) (**H-J**) or control PASMCs treated with 10µM of lactate (**K-M**) for 48 hours. Data are means±SE; n=3-6 subjects/group. *p<0.05, **p<0.01 by Mann Whitney U.

### TOP1 inhibitor indotecan selectively suppresses Akt-mTORC1, reduces proliferation, and induces apoptosis in human PAH PASMCs

To test potential benefits of pharmacological inhibition of TOP1, we evaluated indotecan, a next-generation TOP1 inhibitor recently approved by Food and Drug Administration (FDA) as an anti-cancer treatment. Similar to siRNA TOP1 (Figure 5C-G), indotecan down-regulated Akt-mTORC1 signaling, as evidenced by significant reduction of P-S473 Akt/total Akt and P-S6/total S6 ratios, decreased proliferation, and induced apoptosis in human PAH, but not in control PASMCs (Figure 6), demonstrating that indotecan selectivity eliminates PAH, but not non-diseased, PASMCs and suggesting its potential attractiveness as an anti-remodeling agent for PAH.

**Figure 6.**
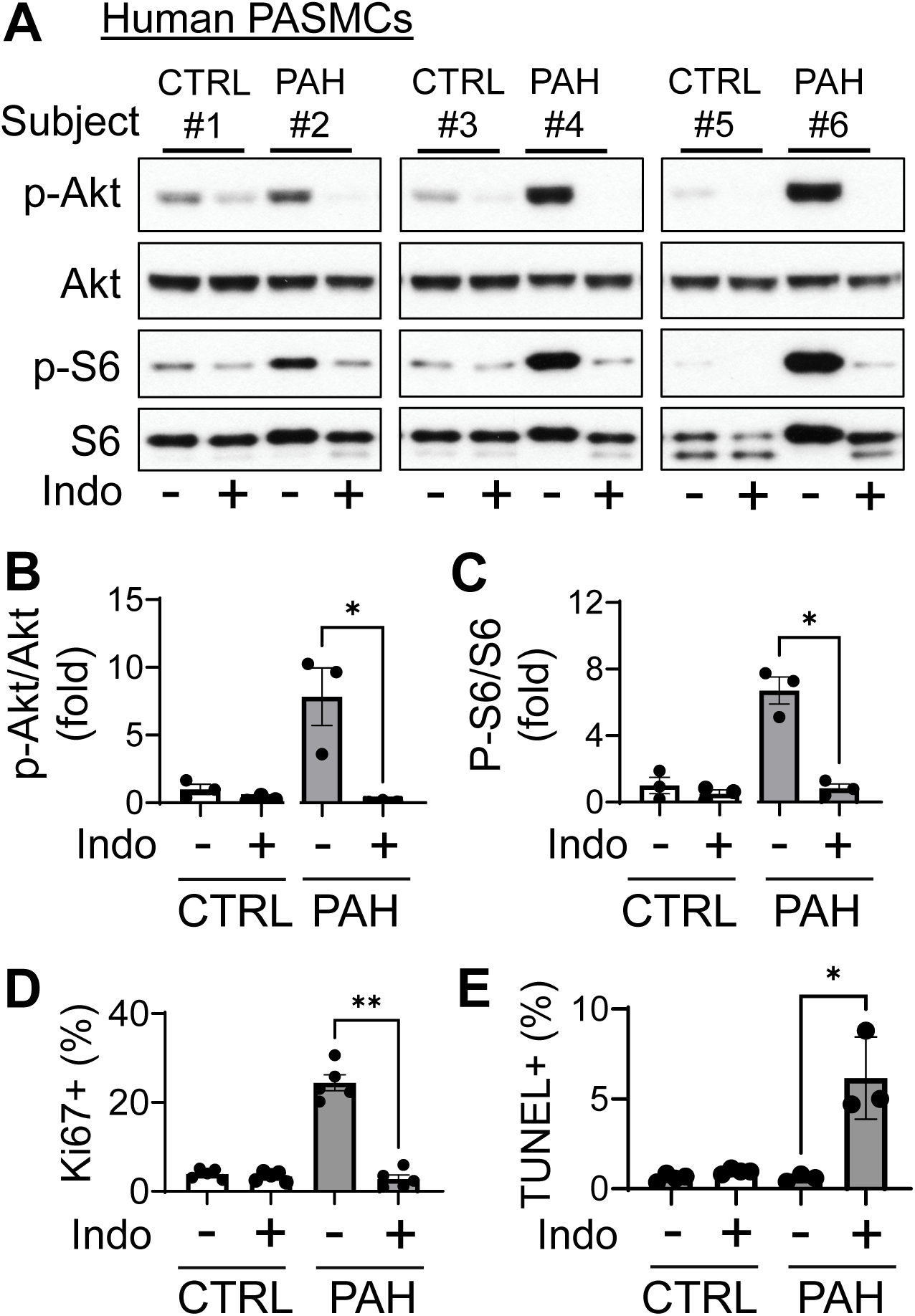
TOP1 inhibitor Indotecan suppresses Akt-mTORC1 and proliferation, induces apoptosis in human PAH, but not control PASMCs. Human control and PAH PASMCs were treated with 0.5 µM indotecan (Indo) or diluent (-) for 48h followed by immunoblot analysis to detect indicated proteins (**A-C**), proliferation (Ki67) (**D**), and apoptosis (*In Situ* Cell Death Detection Kit) (**E**) assays. Data are means±SE, n=3-5 subjects/group, *p<0.05, **p<0.01 by Kruskal-Wallis (Dunn).

### TOP1 inhibitor indotecan normalizes Akt-mTORC1 and reverses severe experimental PH in rats

Next, we evaluated potential benefits of indotecan *in vivo* using severe irreversible SuHx rat model of PH (35, 36). Briefly, rats received single injection of SU5416 and maintained for three weeks under normobaric hypoxia (10% O_2_) followed by two weeks under normoxia. Five weeks after PH induction, rats were randomized into two groups and treated with indotecan or vehicle for next 14 days (Figure 7A). Similar to human PAH (Figure 1G), vehicle-treated SuHx-exposed rats demonstrated significant TOP1 overaccumulation in SMA-positive areas of small remodeled PAs, which was accompanied by significant increase in S473-Akt and S6 phosphorylation, elevated PA MT, sRVP, RV end diastolic pressure (RV EDP), RV contractility (max dP/dT), and Fulton index (RV/(LV+S) (Figure 7B-M). Treatment with indotecan reduced Akt and S6 phosphorylation in small PAs, reversed pulmonary vascular remodeling, sRVP, RV hypertrophy, and normalized RV functional parameters (Figure 6B-M). We observed no significant differences in sLVP and heart rate among groups (Figure S8). These data demonstrate that up-regulation of TOP1 signaling is shared between SuHx rat model of PH and human PAH, and show preclinical benefits of indotecan to reverse pathological Akt-mTORC1 activation, pulmonary vascular remodeling, PH, and RV hypertrophy and dysfunction in severe experimental PH.

**Figure 7.**
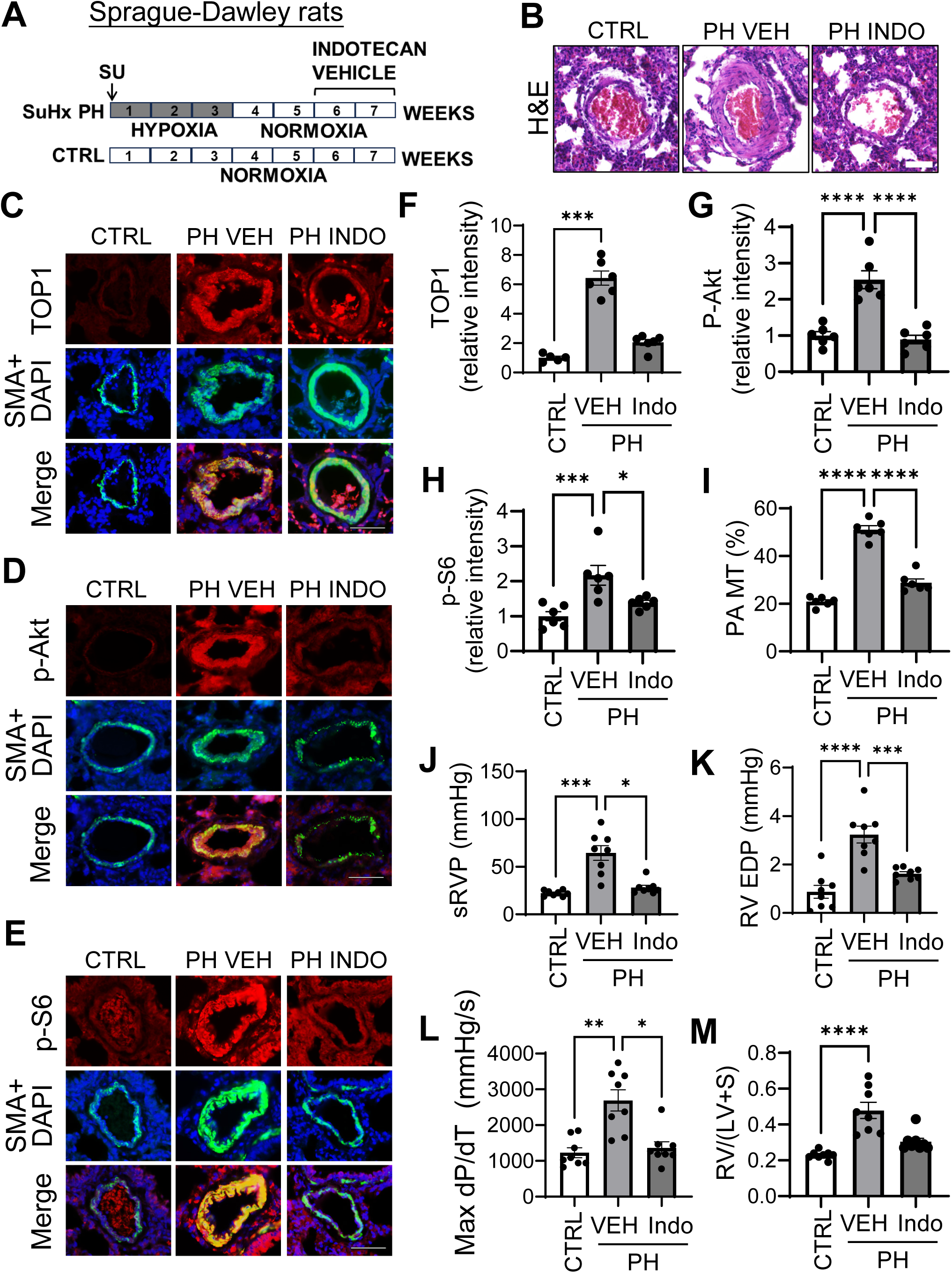
TOP1 inhibitor Indotecan down-regulates Akt-mTORC1 in small PAs, reverses pulmonary vascular remodeling, PH, and improves RV in SuHx rats. **A:** PH was developed in 6-8 weeks-old female Sprague Dawley rats by single SU5416 injection (s.q., 20 mg/kg) followed by three weeks of normobaric hypoxia and 2 weeks of normoxia. Then, animals were randomly assigned to two groups and treated with diluent or Indotecan (2.5 mg/kg/day, i.p., 5 days/week) for next two weeks. Negative controls were normoxia-maintained same-age female rats. Hemodynamic analysis and tissue collection were performed on day 50 of experiment. **B-H:** H&E (**B**) and immunohistochemical analysis of lung tissue sections (**C-H**). Bar=50µm. Images are representative of 5-6 rats/group. **C-E**: Red -TOP1 (**C**), p-S473 Akt (**D**), or P-S6 (**E**), green -SMA, blue -DAPI. **I-M**: PA MT (**I**), sRVP (**J**), EV end-diastolic pressure (RV EDP) (**K**), max dP/dT (**L**), and Fulton index (RV/(LV+S) (**M**) of indicated groups. Data are means±SE, n=6 (**I**) or 8 (**J-M**) rats/group. *p<0.05, **p<0.01, ***p<0.005, ****p<0.0001 by Kruskal-Wallis (Dunn) (**F, I, J**, **L**, **M**) or ANOVA (Tukey) (**G, H, K**).

## DISCUSSION

We here report that over-lactylation and over-accumulation of TOP1 due to LDHA-induced lactate over-production supports PASMC hyper-proliferation, pulmonary vascular remodeling, and PH. Our new findings suggest that LDHA-lactate axis promotes hyper-proliferation and survival of human PAH PASMCs via over-lactylation of TOP1, leading to TOP1 over-accumulation, consequent activation of Akt-mTORC1, PA remodeling, and PH. We also show that the second-generation TOP1 inhibitor, indotecan, normalizes Akt-mTORC1, selectively reduces proliferation and induces apoptosis in human PAH PASMCs, and reverses PA remodeling and severe PH in rats (Figure 8).

**Figure 8.**
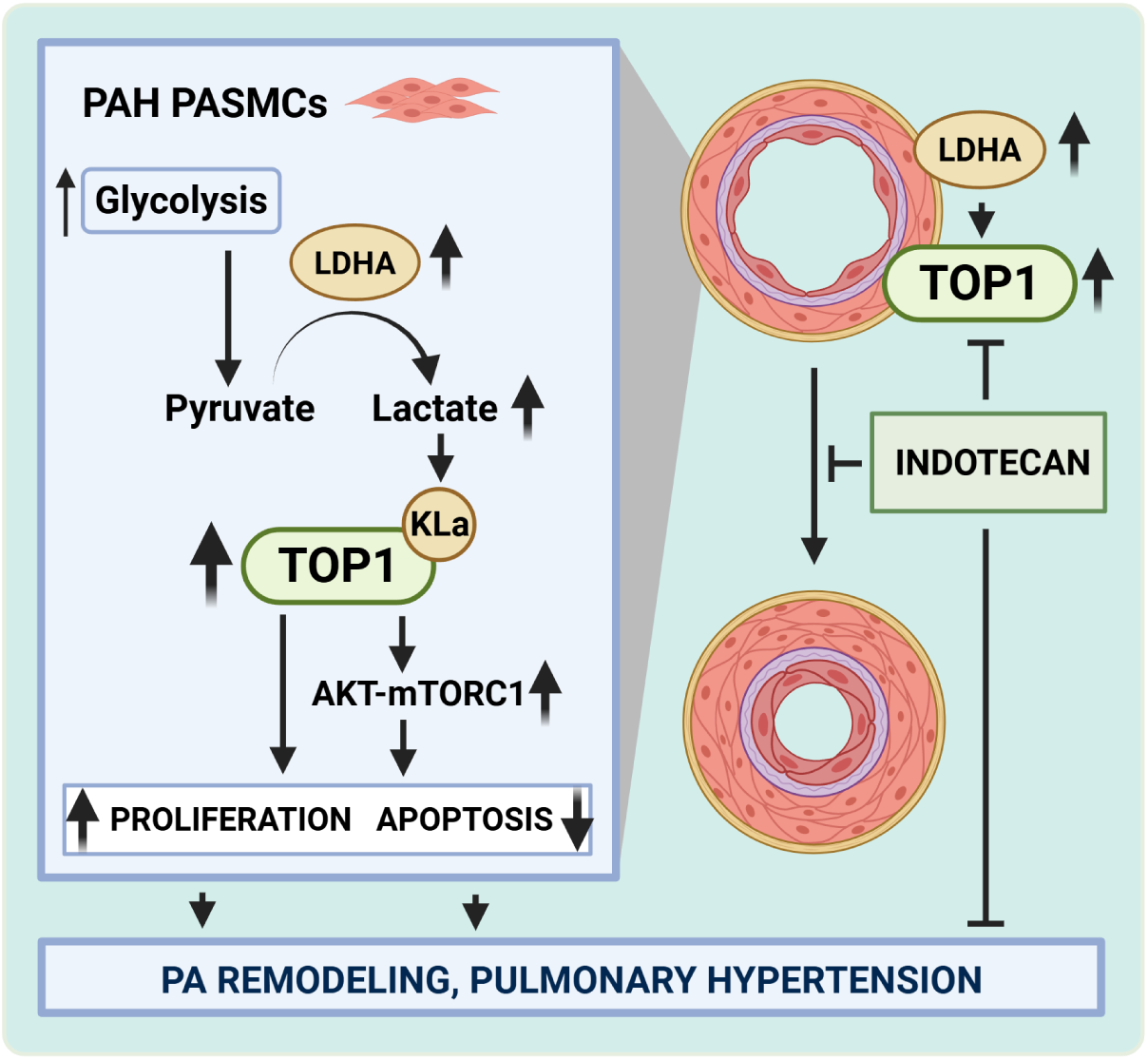
Schematic representation of the findings. LDHA-dependent lactate over-production promotes PASMC proliferation, survival, and pulmonary hypertension via over-lactylation and over-accumulation of TOP1 and consequent Akt-mTORC1 activation.

Metabolic and signaling reprograming of pulmonary vascular cells are key drivers of pulmonary vascular remodeling, an important pathophysiological feature of PAH. The metabolic shift to glycolysis, known as the Warburg effect, supports hyper-proliferative, apoptosis-resistant PASMC phenotype, and PA remodeling in PAH (11, 14, 37, 38), and also leads to over-production of lactate (16, 21). Lactate plays a central role in the self-sufficient metabolism, proliferation, and sustainability of glycolytic cancer cells (39, 40), and promotes PASMC proliferation (18–21), suggestive of its potential disease-modifying role in human PAH. We found that LDHA, an enzyme which converts pyruvate to lactate, is over-expressed in PASMCs in small muscular PAs in PAH lungs, driving lactate over-production, consequent increase in TOP1 over-lactylation, cell proliferation and apoptosis resistance.

Our findings identify DNA topoisomerase TOP1 as a mechanistic link among glycolysis, Akt-mTORC1 activation, and PAH PASMC proliferation and survival, providing important insight into the interplay between metabolic and molecular abnormalities in PAH pulmonary vasculature. Selective inhibitors of TOP1 are already in clinical use to treat cancer (41, 42), making it a translationally promising over-lactylated protein identified by us in this study. However, it is important to note that TOP1 is only one of 12 proteins identified as over-lactylated in PASMCs from human PAH lungs by unbiased proteomic analysis. Another identified protein, tenascin-C (TNC), is a well-described pro-remodeling factor, over-accumulation of which in PAH pulmonary vasculature supports proliferation and migration of PASMCs (43, 44). The rest of the proteins have not been studied in PAH settings, but have been linked to cancer and/or cardiovascular disease (45–55), and we can’t exclude their potential involvement in PASMC remodeling and PAH.

Lactylation of non-histone proteins regulates their behavior by modulating protein abundance, activity, or localization (56). We found that over-lactylation of TOP1 was accompanied by its over-accumulation in PASMCs in small muscular PAs from human PAH and experimental PH lungs. Intriguingly, a highly probable linkage of TOP1 with the PAH network were in the Akt-mTOR, p53, and WNT-β catenin pathways, critical regulators of cell proliferation, apoptosis, and remodeling in PAH (9, 34, 57–60). Further validation revealed that TOP1 and LDHA-lactate positively regulate Akt-mTORC1 in human PASMCs. Together with published studies (34, 58, 59), our data establishes a new mechanistic links from the glycolysis-and LDHA-driven lactate overproduction, via lactylation and overaccumulation of TOP1, to the up-regulation of Akt-mTORC1, hyper-proliferation, and apoptosis resistance of PASMCs in PAH. We didn’t observe LDHA-lactate-TOP1-dependent modulation of p53 and/or β-catenin protein levels. However, in cancer cells, p53 serves as an upstream regulator of TOP1 activation by DNA damage, and β-catenin uses DNA topoisomerase Topo IIα as a transcriptional co-activator (61, 62). Thus, we can’t exclude that p53 and WNT-β-catenin act upstream of TOP1 or that TOP1 regulates these proteins via different signaling mechanisms.

The remodeling-focused therapies for patients with PAH are limited to sotatercept (63). Our findings establish LDHA-TOP1 axis as a promising candidate target pathway to treat established PAH. Our observations that LDHA inhibitor oxamate down-regulates TOP1 and Akt, reverses pulmonary vascular remodeling, PH, and RV hypertrophy in mice with SuHx PH are in good agreement with a recent report showing that LDHA inhibitor GSK2837808A reduces Akt phosphorylation, attenuates PA remodeling and experimental PH in mice and rats (18). The LDHA inhibitors, including GSK2837808A, are currently at the preclinical development stage as anti-cancer agents, with one (CHK-336) at phase I trials for healthy volunteers (ClinicalTrials.gov Identifier: NCT05367661), making it challenging to translate these findings to patient care.

The most clinically advanced to date are TOP1 inhibitors, the first generation of which, topotecan and irinotecan, have been widely used as anticancer drugs for more than 20 years (41, 64). Recently, a new generation TOP1 inhibitor, indotecan (LMP744), was approved by the FDA for the treatment of patients with malignant glioma. Our preclinical study demonstrated that indotecan selectively suppresses Akt-mTORC1 signaling, eliminates “diseased” human PAH PASMCs without affecting non-diseased cells *in vitro,* and reverses SuHx-induced pulmonary vascular remodeling, PH, and restores RV morphology and function *in vivo*, providing clear evidence of potential benefits of using indotecan to treat PAH.

There are several limitations in our study, including a small sample size of human specimens-based experiments. PAH is rare disease, and the availability of human tissue specimens and primary cells of early passage is limited. However, the important role of the LDHA-lactate-TOP1 axis in hyper-proliferative PASMC phenotype, PA remodeling, and overall PH is well supported by a siRNA-and an inhibitors-based approach as well as an analysis of mice with SM-specific Ldha loss. Another limitation is that only female rats were preclinically evaluated in the rat SuHx model of PH as we didn’t observe sex-related differences between Ldha-/-^SMC^ male and female animals. In addition, the indotecan treatment was administered to the animals as a mono-treatment, and its combination with currently used FDA-approved therapies remains to be evaluated. However, given our findings in the PASMCs from PAH subjects and transgenic mice, further testing of indotecan as an add-on therapy to reverse PAH is worthy of further investigation. Lastly, we can’t exclude that lactate plays a role in other groups of PH and/or associated diseases. Histone lactylation plays an important role in experimental hypoxia-induced PASMC proliferation, senescence, and PH, and topotecan was recently shown to prevent development of hypoxic PH in rats (24, 65–68), suggestive of potential involvement in group 3 PH associated with lung diseases and/or hypoxemia. Elevated lactate levels are linked with adverse clinical outcomes in COPD (69) and are recently reported as a marker of reduced exercise capacity in patients with heart failure with preserved ejection fraction (70), opening new exciting avenues for further studies.

In conclusion, our findings identify LDHA-lactate-driven TOP1 signaling as a new molecular pathway linking metabolic abnormalities with pulmonary vascular remodeling and PAH. Our data also suggest that TOP1 could be considered as a promising remodeling-focused molecular target, and TOP1 inhibitor indotecan as a potential disease-modifying therapy to treat PAH.

## MATERIALS AND METHODS

(expanded in the Online Data Supplement)

### Human tissues and cell cultures

The human lung tissues and early-passage (3–8) human PASMCs, PAECs, and PAAFs were isolated from small (<1.5 mm outer diameter) PAs of non-diseased subjects (postmortem tissue acquisition) and patients with PAH (lung explants after transplantation procedure) were provided by the Pulmonary Hypertension Breakthrough Initiative (PHBI), University of Pittsburgh VMI Cell Processing Core, University of California at San Francisco transplant program, and University of California at Davis Lung Center Pulmonary Vascular Disease Program under approved protocols in accordance with Institutional Review Boards and the Committee for Oversight of Research and Clinical Training Involving Decedents (see Table S1 for human subjects characteristics). Cell isolation, characterization, and maintenance were performed under the rigorous and well-established protocols adopted by the PHBI (14, 26, 71, 72). Primary human vascular cells of the same passage from a minimum of three subjects were used in each experiment. Cells were maintained in complete Smooth Muscle Cell Growth Medium 2 (PASMCs), complete Fibroblast Growth Medium (PAAFs), or complete Endothelial Cell Growth Medium MV2 (PAEC) (basal medium + supplement pack) and Antibiotic-Antimycotic. Before experiments, cells were maintained in the basal media supplemented with 5% fetal bovine serum (FBS) and serum-deprived for 24-48 hours in basal media supplemented with 0.1% bovine serum albumin (BSA) (PASMCs, PAAFs) or for 24 hours in basal media supplemented with 0.5% FBS (PAECs) if not stated otherwise.

### Animals

All animal procedures were performed under the protocols approved by the Animal Care and Use Committee of the University of California, Davis. Ldha^-/-SMC^ mice were generated by crossing Ldha^flox^ (B6(Cg)-Ldha^tm1c(EUCOMM)Wtsi^/DatsJ, Cat#030112) mice with SM22-Cre mice (B6.Cg-Tg(Tagln-cre)1Her/J, Cat#017491) from The Jackson Laboratory (Sacramento, CA). Animal genotyping was performed in accordance with manufacturer’s protocol. Male and female Ldha^-/-SMC^, wild type mice of the same background (C57BL/6J) (The Jackson Laboratory Cat #000664), and female Sprague-Dawley rats (Charles River Laboratories, Hollister, CA, Cat#001) were used in experiments. PH in mice was induced by SuHx exposure as described in (25, 26, 36). Oxamate (750 mg/kg, i.p.) or vehicle (saline, i.p.) were administrated daily on days 22-35 of experiment. Blinded hemodynamic analysis and tissue harvest were performed on days 21 and 35. SuHx PH in rats was induced as described previously (35, 59). Indotecan or vehicle were administrated on days 36-49 of the experiment (2.5 mg/kg/day, i.p., 5 days/week). Negative controls were normoxia-maintained same-age female rats. At day 50, terminal hemodynamic analysis was performed as described in (26, 35, 59, 73), morphological and immunohistochemical analyses were performed, the Fulton index was calculated by the RV/(LV+Septum) weight ratio as described in (14, 25, 26, 35, 36, 59, 74). Animals’ randomization and blinded analysis were performed when possible.

### Immunohistochemical, immunoblot, RT PCR, proliferation, apoptosis analyses, and transfection

were performed as described previously (14, 25, 26, 36) (14, 25, 26, 36).

### Mass Spectrometry analysis

was performed by MS Bioworks (Ann Arbor, MI) as described previously (25).

### Statistical analysis

was performed using StatView and GraphPad Prism software. Statistically significant differences among groups for datasets with n<6/group or n≥6/group without normal distribution were assessed with Mann-Whitney U (2 groups) and Kruskal-Wallis (Dunn correction) tests (≥3 groups). Normally distributed data with n≥6/group was analyzed using t-test (2 groups) or ANOVA (Tukey’s correction) (≥3 groups). Statistical significance was defined as p<0.05. No animals were excluded from the analysis.

